# A Connectome of the Adult *Drosophila* Central Brain

**DOI:** 10.1101/2020.01.21.911859

**Authors:** C. Shan Xu, Michal Januszewski, Zhiyuan Lu, Shin-ya Takemura, Kenneth J. Hayworth, Gary Huang, Kazunori Shinomiya, Jeremy Maitin-Shepard, David Ackerman, Stuart Berg, Tim Blakely, John Bogovic, Jody Clements, Tom Dolafi, Philip Hubbard, Dagmar Kainmueller, William Katz, Takashi Kawase, Khaled A. Khairy, Laramie Leavitt, Peter H. Li, Larry Lindsey, Nicole Neubarth, Donald J. Olbris, Hideo Otsuna, Eric T. Troutman, Lowell Umayam, Ting Zhao, Masayoshi Ito, Jens Goldammer, Tanya Wolff, Robert Svirskas, Philipp Schlegel, Erika R. Neace, Christopher J. Knecht, Chelsea X. Alvarado, Dennis A. Bailey, Samantha Ballinger, Jolanta A Borycz, Brandon S. Canino, Natasha Cheatham, Michael Cook, Marisa Dreher, Octave Duclos, Bryon Eubanks, Kelli Fairbanks, Samantha Finley, Nora Forknall, Audrey Francis, Gary Patrick Hopkins, Emily M. Joyce, SungJin Kim, Nicole A. Kirk, Julie Kovalyak, Shirley A. Lauchie, Alanna Lohff, Charli Maldonado, Emily A. Manley, Sari McLin, Caroline Mooney, Miatta Ndama, Omotara Ogundeyi, Nneoma Okeoma, Christopher Ordish, Nicholas Padilla, Christopher Patrick, Tyler Paterson, Elliott E. Phillips, Emily M. Phillips, Neha Rampally, Caitlin Ribeiro, Madelaine K Robertson, Jon Thomson Rymer, Sean M. Ryan, Megan Sammons, Anne K. Scott, Ashley L. Scott, Aya Shinomiya, Claire Smith, Kelsey Smith, Natalie L. Smith, Margaret A. Sobeski, Alia Suleiman, Jackie Swift, Satoko Takemura, Iris Talebi, Dorota Tarnogorska, Emily Tenshaw, Temour Tokhi, John J. Walsh, Tansy Yang, Jane Anne Horne, Feng Li, Ruchi Parekh, Patricia K. Rivlin, Vivek Jayaraman, Kei Ito, Stephan Saalfeld, Reed George, Ian Meinertzhagen, Gerald M. Rubin, Harald F. Hess, Louis K. Scheffer, Viren Jain, Stephen M. Plaza

## Abstract

The neural circuits responsible for behavior remain largely unknown. Previous efforts have reconstructed the complete circuits of small animals, with hundreds of neurons, and selected circuits for larger animals. Here we (the FlyEM project at Janelia and collaborators at Google) summarize new methods and present the complete circuitry of a large fraction of the brain of a much more complex animal, the fruit fly *Drosophila melanogaster*. Improved methods include new procedures to prepare, image, align, segment, find synapses, and proofread such large data sets; new methods that define cell types based on connectivity in addition to morphology; and new methods to simplify access to a large and evolving data set. From the resulting data we derive a better definition of computational compartments and their connections; an exhaustive atlas of cell examples and types, many of them novel; detailed circuits for most of the central brain; and exploration of the statistics and structure of different brain compartments, and the brain as a whole. We make the data public, with a web site and resources specifically designed to make it easy to explore, for all levels of expertise from the expert to the merely curious. The public availability of these data, and the simplified means to access it, dramatically reduces the effort needed to answer typical circuit questions, such as the identity of upstream and downstream neural partners, the circuitry of brain regions, and to link the neurons defined by our analysis with genetic reagents that can be used to study their functions.

Note: In the next few weeks, we will release a series of papers with more involved discussions. One paper will detail the hemibrain reconstruction with more extensive analysis and interpretation made possible by this dense connectome. Another paper will explore the central complex, a brain region involved in navigation, motor control, and sleep. A final paper will present insights from the mushroom body, a center of multimodal associative learning in the fly brain.

## 1 Introduction

The connectome we present here is a dense reconstruction of a portion of the central brain (referred to as the hemibrain) of the fruit fly, *Drosophila melanogaster*, as shown in Fig 1. This region was chosen since it contains all the circuits of the central brain (assuming lateral symmetry), and in particular contains circuits critical to unlocking mysteries involving associative learning in the mushroom body, navigation and sleep in the central complex, and circadian rhythms among clock circuits. The largest dense reconstruction to date, it contains around 25,000 neurons, most of which were rigorously clustered and named, with about 20 million chemical synapses between them, plus portions of many other neurons truncated by the boundary of the data set (details in Fig. 1 below). Each neuron is documented at many levels - the detailed voxels that constitute it, a simplified skeleton, and the synaptic partners and location of most synapses.

**Figure 1:**
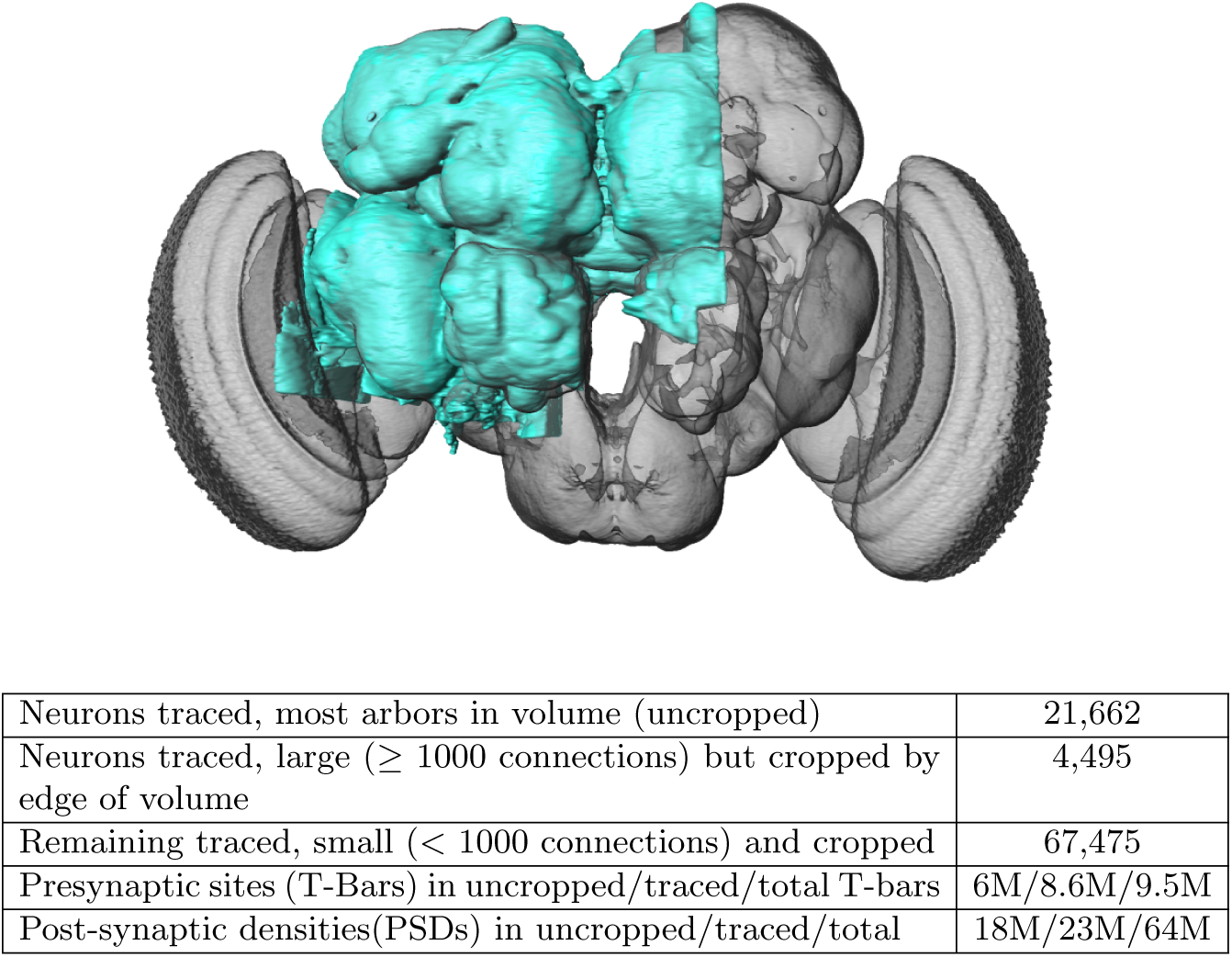
The hemibrain and some basic statistics. The highlighted area shows the portion of the central brain that was imaged and reconstructed, superimposed on a grayscale representation of the whole *Drosophila* brain. For the table, a neuron is traced if all its main branches within the volume are reconstructed. A neuron is considered uncropped if most arbors (though perhaps not the soma) are contained in the volume. Otherwise it is cropped. Note: 1) our definition of cropped is somewhat subjective, 2) the usefulness of a cropped neuron depends on the application, and 3) some small fragments are known to be distinct neurons. For simplicity, we will often state the hemibrain contains 25K neurons.

Producing this data set required advances in sample preparation, imaging, image alignment, machine segmentation of cells, synapse detection, data storage, proofreading software, and protocols to arbitrate each decision. A number

These data describe whole brain properties and circuits, as well as contain new methods of cell type classification based on connectivity. Computational compartments are now more carefully defined, we can identify actual synaptic circuits, and each neuron is annotated by name and putative cell type, making this the first complete census of neuropils, tracts, cells, and connections in this portion of the brain. We compare the statistics and structure of different brain regions, and the brain as a whole, without the confounds from studying different circuitry in different animals.

All data are publicly available through web interfaces. This includes a browser interface, NeuPrint[2], designed so that any interested user, even without specific training, can query the hemibrain connectome. NeuPrint can query the connectivity, partners, connection strengths and morphologies of any specified neurons, thus making identification of upstream and downstream partners orders of magnitude easier than through existing genetic methods. In addition, for those who are willing to program, the full data set - the gray scale voxels, the segmentation and proofreading results, skeletons, and graph model of connectivity, are also available through publicly accessible application program interfaces (APIs).

This effort differs from previous EM reconstructions in its social and collaborative aspects. Previous reconstructions, whether dense in small EM volumes(such as [3][4]), or sparse in larger volumes (such as [5] or [6]), have concentrated on reconstruction of specific circuits to answer specific questions. When the same EM volume is used for many such efforts, as occurred in the *Drosophila* larva and the full adult fly brain, this leads to an overall reconstruction that is the union of many individual efforts[7]. The result is inconsistent coverage of the brain, with some areas well reconstructed and others missing entirely. In contrast, here we have analyzed the whole volume, not just the subsets interesting to specific groups of researchers with the time, energy and expertise to tackle EM reconstruction. We are making this data available without restriction, with only the requirement to cite the source. This allows the benefits of known circuits and connectivity to accrue to the field as a whole, a much larger audience than those with expertise in EM reconstruction. This is analogous to progress in genomics, which transitioned from individual groups studying subsets of genes, to publicly available genomes that can be queried for information about genes of choice[8].

One major benefit to this effort is to facilitate research into the circuits of the fly brain. A common question among researchers, for example, is the identity of upstream and downstream partners of specific neurons. Previously this could only be addressed by genetic methods such as trans-Tango[9] or by sparse tracing in previously obtained EM volumes[6]. These methods are technically difficult, require specialized expertise, and are time consuming, often taking months of effort. Now, for any circuits contained in our volume, any researcher can obtain the same answers in seconds by querying a publicly available database.

Another major benefit of dense reconstruction is its exhaustive nature. Genetic methods such as stochastic labeling may miss cell types, and counts of cells of a given type are dependent on expression levels, which are always uncertain. Previous dense reconstructions have demonstrated that existing cell type catalogs are incomplete, even in well-covered regions[4]. In our hemibrain sample, we have identified all the cells within the reconstructed volume, thus providing a complete and unbiased census of all cell types in the fly central brain (at least in this single female animal), and a precise count of the instances of each type. Another scientific benefit is analysis without the uncertainty of pooling data obtained with different animals. The detailed circuitry of the fly brain is known to depend on nutritional history, age, and circadian rhythm. Here these factors are held constant, as are the experimental methods, facilitating comparison of different fly brain regions in this single animal. Evaluating stereotypy across animals will require additional connectomes.

### 1.1 What is included

Fig. 2 shows the hierarchy of the named brain regions that are included in the hemibrain. Table 1 shows the primary regions that are at least 50% included in the hemibrain sample, their approximate size, and their completion percentage. Our names for brain regions follow the conventions of [1] with the addition of ‘(L)’ or ‘(R)’ to indicate whether the region (most of which occur on both sides of the fly) has its cell bodies in the left or right, respectively. The mushroom body[10][11] and central complex[12] are further divided into finer compartments.

**Figure 2:**
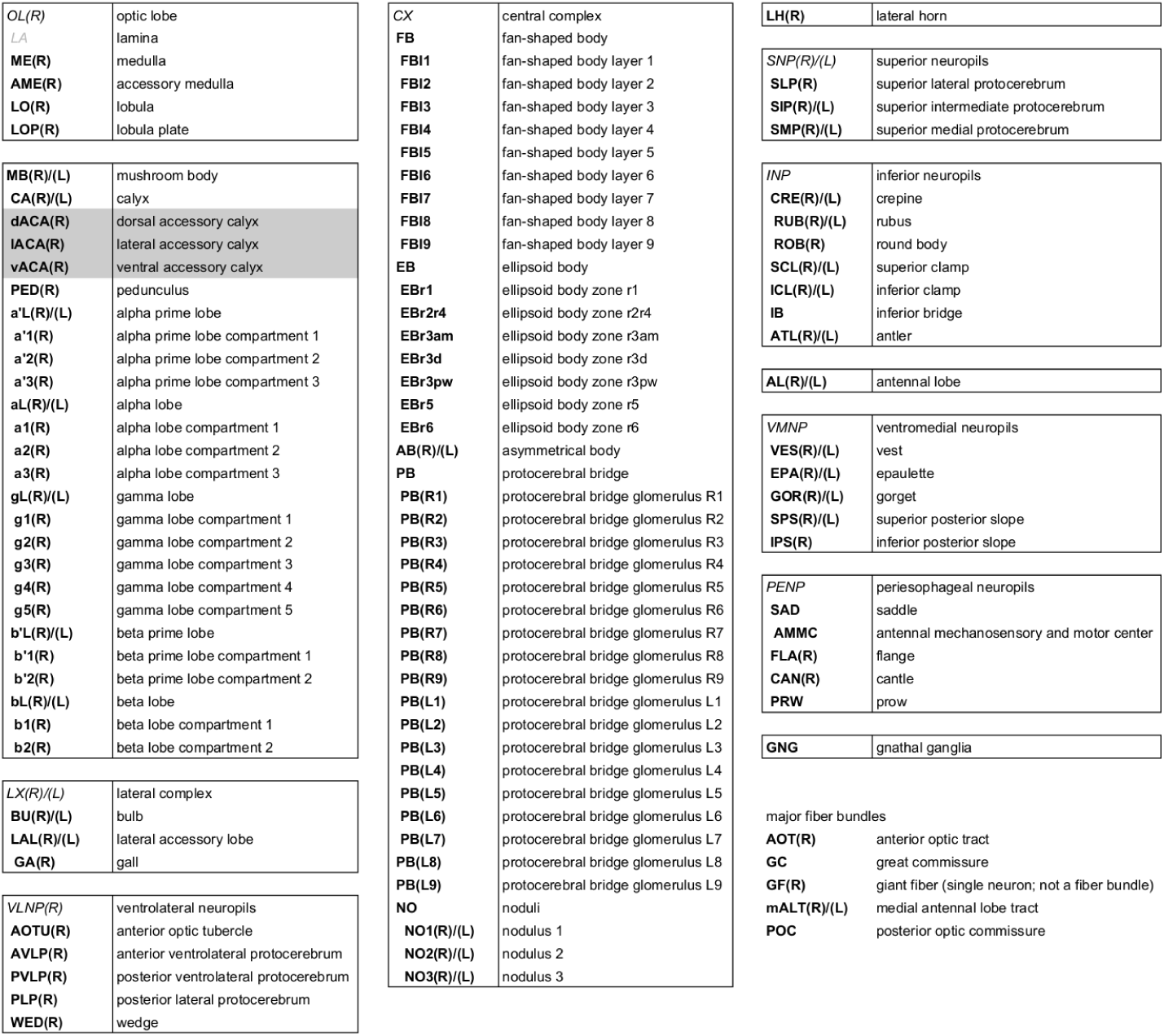
Brain regions contained and defined in the hemibrain, following the naming conventions of [1] with the addition of (R) and (L) to specify the side of the soma for that region. Gray *italics* indicate master regions not explicitly defined in the hemibrain. Region LA is not included in the volume. The regions are hierarchical, with the more indented regions subsets of the less indented regions. The only exceptions are dACA, lACA, and vACA which are considered part of the mushroom body but are not contained in the master region MB. of new tests for estimating the completeness and accuracy were required and therefore developed, in order to verify the correctness of the connectome.

**Table 1:** Regions that are at least 50% included in the hemibrain, sorted by completion percentage. ‘%inV’ is the approximate percentage of the region included in the hemibrain volume. ‘T-bars’ gives a rough estimate of the size of the region. ‘comp%’ is the fraction of the PSDs contained in the brain region for which both the PSD, and the corresponding T-bar, are in neurons marked as ‘Traced’.

| Name | %inV | T-bars | comp% | Name | %inV | T-bars | comp% |
| --- | --- | --- | --- | --- | --- | --- | --- |
| PED(R) | 100% | 54805 | 85% | aL(R) | 100% | 95375 | 84% |
| b'L(R) | 100% | 67695 | 83% | bL(R) | 100% | 71112 | 83% |
| gL(R) | 100% | 176785 | 83% | a'L(R) | 100% | 39091 | 82% |
| EB | 100% | 164286 | 81% | bL(L) | 56% | 58799 | 80% |
| NO | 100% | 36722 | 79% | b'L(L) | 88% | 57802 | 78% |
| gL(L) | 55% | 133256 | 76% | CA(R) | 100% | 69515 | 73% |
| AB(R) | 100% | 2734 | 64% | aL(L) | 51% | 44803 | 62% |
| FB | 100% | 451040 | 61% | AL(R) | 83% | 501007 | 58% |
| AB(L) | 100% | 572 | 57% | PB | 100% | 46557 | 55% |
| AME(R) | 100% | 6045 | 47% | BU(R) | 100% | 9381 | 46% |
| CRE(R) | 100% | 137946 | 39% | AOTU(R) | 100% | 92579 | 37% |
| LAL(R) | 100% | 234398 | 36% | SMP(R) | 100% | 510943 | 33% |
| PVLP(R) | 100% | 475228 | 29% | ATL(R) | 100% | 25472 | 28% |
| SPS(R) | 100% | 253821 | 28% | ATL(L) | 100% | 28153 | 28% |
| VES(R) | 84% | 157171 | 27% | IB | 100% | 200447 | 27% |
| CRE(L) | 90% | 130498 | 27% | SIP(R) | 100% | 187494 | 26% |
| BU(L) | 52% | 7014 | 26% | GOR(R) | 100% | 27140 | 25% |
| WED(R) | 100% | 232901 | 24% | SMP(L) | 100% | 460793 | 24% |
| EPA(R) | 100% | 31439 | 24% | PLP(R) | 100% | 429106 | 24% |
| AVLP(R) | 100% | 630542 | 22% | ICL(R) | 100% | 202550 | 22% |
| SLP(R) | 100% | 475903 | 21% | LO(R) | 64% | 855261 | 21% |
| SCL(R) | 100% | 187674 | 21% | GOR(L) | 60% | 19558 | 20% |
| LH(R) | 100% | 231667 | 18% | CAN(R) | 68% | 6513 | 15% |

### 1.2 Differences from connectomes of vertebrates

Most textbooks, such as *Principles of Neural Science*[13], define the operation of the mammalian nervous system with, at most, only passing reference to invertebrate brains. Fly (or other insect) nervous systems differ from those of mammals in several respects. Some main differences include:

- The vast majority of synapses are polyadic, meaning each synapse structure consists of one release site and several adjacent cells with receptors. An element, typically called a T-bar, marks the site of transmitter release into the cleft between cells. This site typically abuts several other cells, where a post-synaptic density (PSD) marks the receptor location.
- Most neurites are not purely axonic or dendritic, but have mixtures of both pre- and post-synaptic partners. Within a single brain region, however, most neurites can be described as mostly dendritic or mostly axonic.
- Unlike synapses in mammals, EM imagery (at least as we have analyzed it here) does not provide obvious information about whether a synapse is excitatory or inhibitory.
- The soma or cell body of each fly neuron resides on the periphery of the brain, mostly disjoint from the main processes innervating the volume. There are no synapses directly on the soma. The neuronal process between the soma and the first branch point is called cell body fiber(CBF) and is not involved in the synaptic transmission of information.
- Synapse sizes are much more uniform than those of mammals. Stronger connections are made by synapses in parallel, not by larger synapses, as in vertebrates. In this paper we will refer to the ‘strength’ of a connection as the synapse count, though we recognize that this is not a true measure of the coupling strength, since we do not know the relative activity and strength of the synapses.
- The brain is small, about 250 *µ*m per side, and lacks myelinated axons. The whole brain has roughly the same size as the dendritic arbor of a single pyramidal neuron in the mammalian cortex.
- Graded connections (as opposed to those of spiking neurons) lack anatomical distinction but are not uncommon. Some neurons even switch between graded and spiking operation[14].

## 2 Connectome Reconstruction

Producing a connectome consisting of reconstructed neuron morphologies and the chemical synapses between them required several steps. The first step, preparing a fly brain and imaging half of its center, produced a dataset consisting of 26 teravoxels of data, each with 8 bits of information. We applied numerous machine learning algorithms and over 50 person-years of proofreading effort over 2 calendar years to extract a variety of more compact and useful representations, such as neuron skeletons, synapse locations, and connectivity graphs. These are both more useful and much smaller than the raw grayscale data - for example, the connectivity can be reasonably summarized by a graph with 25,000 nodes and 3 million edges. Even with connections broken down by brain region, such a graph takes only 26 MB, roughly a million fold reduction in data size.

Many of the supporting methods for this reconstruction have been recently published by the members of our team. Here we briefly survey each major area, with more details found in the companion papers. Major advances include:

- New methods to fix and stain the sample, preparing a whole fly brain with well-preserved subcellular detail particularly suitable for machine analysis.
- Methods to enable the largest yet EM data collection using Focused Ion Beam Scanning Electron Microscopy (FIB-SEM) technology, resulting in isotropic data with few artifacts, a combination that significantly speeds up reconstruction.
- A coarse-to-fine, automated flood-filling network segmentation pipeline applied to image data normalized with cycle-consistent generative adversarial networks, and an aggressive automated agglomeration regime enabled by advances in proofreading.
- A new hybrid synapse prediction method, using two differing underlying technologies, for accurate synapse prediction across the volume.
- New top-down proofreading methods that leverage visualization and machine learning to achieve orders of magnitude faster reconstruction compared to previous approaches in the fly’s brain.

These are summarized below.

### 2.1 Image stack collection

The first step, fixing and staining the specimen, has been accomplished taking advantage of three new developments. These improved methods allow us to fix and stain a full fly brain and still recover neurons as round profiles with darkly stained synapses, suitable for machine segmentation and automatic synapse detection. Starting with a five day old female of wild-type Canton S strain G1 x w1118, we use a custom-made jig to microdissect the central nervous system, which was then fixed and embedded in Epon, an epoxy resin. We then enhanced the electron contrast by staining with heavy metals, and progressively lowered the temperature during dehydration of the sample. Collectively these methods optimize morphological preservation, allow full-brain preparation without distortion (unlike fast freezing methods), and provide increased staining intensity that allows for faster FIB-SEM imaging[15].

The hemibrain sample is roughly 250 x 250 x 250 *µ*m^3^, larger than we can FIB-SEM without introducing milling artifacts. Therefore we subdivided our plastic-embedded samples into 20 *µ*m thick slabs, both to avoid artifacts and allow imaging in parallel for increased throughput. To be effective, the cut surfaces of the slabs must be smooth at the ultrastructural level and have only minimal material loss. Specifically, for connectomic research, all long-distance processes must remain traceable across sequential slabs. We used an improved version of our previously published ‘hot-knife’ ultrathick sectioning procedure[16] which uses a heated, oil-lubricated diamond knife, to section the *Drosophila* brain into 37 sagittal slabs of 20 *µ*m thickness with an estimated material loss between consecutive slabs of ~30 nm - small enough to allow tracing of long-distance neurites. Each slab was re-embedded, mounted, and trimmed, then examined with a 3D microCT scan to check for sample quality and establish a scale factor for the Z cutting by FIB. The resulting slabs were FIB-SEM imaged separately (often in parallel, for increased throughput) and the resulting volume datasets were stitched together computationally.

Connectome studies come with clearly defined resolution requirements - the finest neurites must be traceable by humans and should be reliably segmented by automated algorithms[17]. In *Drosophila* connectomic research, the very finest neural processes can be as little as to 15 nm[18]. The fundamental biological dimension determines the minimum isotropic resolution requirements for tracing neural circuits. To meet the high isotropic resolution and large volume imaging demand, we chose the FIB-SEM imaging platform, which offers high isotropic resolution (< 10 nm in x, y, and z), minimal artifacts, and robust image alignment. The high-resolution and isotropic dataset possible with FIB-SEM has substantially sped up the *Drosophila* connectome pipeline. The automated segmentation generates fewer errors due to the higher quality of raw images, the small number of artifacts, and the isotropic resolution. Manual proofreading and correction are also easier and faster with isotropic imaging with minimal defects.

However, when we began, and even now, deficiencies in imaging speed and system reliability of any commercial FIB-SEM system capped the maximum possible image volume to less than 0.01% of a full fly brain. To remedy this, we redesigned the entire control system from the ground up, improved the image speed more than 10x, and created innovative solutions addressing all known failure modes, which thereby expanded the practical imaging volume of conventional FIB-SEM by more than four orders of magnitude from 10^3^*µ*m^3^ to 3 10^7^*µ*m^3^, while maintaining an isotropic resolution of 8 x 8 x 8 nm^3^ voxels[19][20]. In order to overcome the aberration of a large field of view (up to 300 *µ*m wide), we developed a novel tiling approach without sample stage movement, in which the imaging parameters of each tile are individually optimized through an in-line auto focus routine without overhead[21]. After numerous improvements, we have transformed the conventional FIB-SEM from a laboratory tool that is unreliable for more than a few days to a robust volume EM imaging platform with effective long-term reliability, able to perform years of continuous imaging without defects in the final image stack. Imaging time is now the main impediment to even larger volumes, rather than FIB-SEM reliability.

In our study here, the *Drosophila* hemibrain, thirteen consecutive hot-knifed slabs were imaged using two customized enhanced FIB-SEM systems, in which an FEI Magnum FIB column was mounted at 90*^◦^* onto a Zeiss Merlin SEM. After data collection, streaking artifacts along the FIB milling direction generated from secondary electrons were computationally removed using a mask in the frequency domain. The image stacks were then aligned using a customized version of the software platform developed for serial section transmission electron microscopy [6][22], followed by binning along z-axis to form the final 8 x 8 x 8 nm^3^ voxel datasets. Milling thickness variations in the aligned series were compensated using a modified version of the method described by Hanslovsky et al.[23], with the absolute scale calibrated by reference to the MicroCT images.

The 20 *µ*m slabs generated by the hot-knife sectioning are re-imbedded in larger plastic tabs prior to FIB-SEM imaging. To correct for the warping of the slab that can occur in this process, methods adapted from Kainmueller[24] were used to find the tissue-plastic interface and flatten each slab.

The series of flattened slabs was then stitched using a custom method for large scale deformable registration to account for deformations introduced during sectioning, imaging, embedding, and alignment (Saalfeld et al. in prep). These volumes were then contrast adjusted using slice-wise contrast limited adaptive histogram equalization (CLAHE)[25], and converted into a versioned database(Distributed, Versioned, Image-oriented Database, or DVID), which formed the raw data for the reconstruction, illustrated in Fig 3.

**Figure 3:**
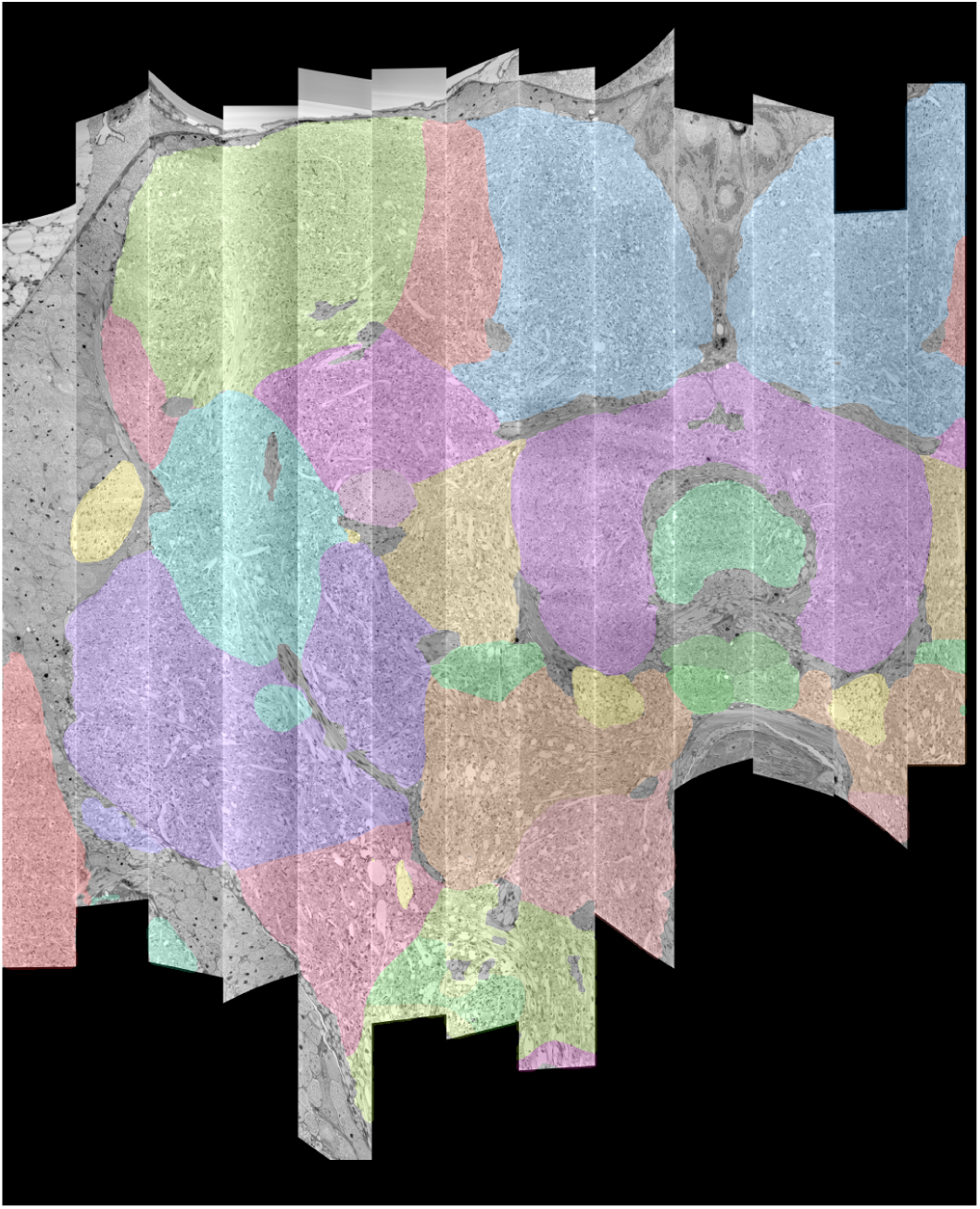
The 13 slabs of the hemibrain, each flattened and co-aligned. Colors are arbitrary and added to the monochrome data to indicate the defined brain regions

### 2.2 Automated Segmentation

Computational reconstruction of the image data was performed using flood-filling networks (FFNs) trained on roughly five-billion voxels of volumetric ground truth contained in two tabs of the hemibrain dataset[17]. Initially, the FFNs generalized poorly to other tabs of the hemibrain whose image content had a different appearance. Therefore we adjusted the image content to be more uniform using cycle-consistent generative adversarial networks (CycleGANs)[26]. Specifically, generator networks were trained to alter image content such that a second discriminator network was unable to distinguish between image patches sampled from, for example, a tab that contained volumetric training data versus a tab that did not. A cycle-consistency constraint was used to ensure the image transformations preserved ultrastructural detail.

FFNs were applied to the CycleGAN-normalized data in a coarse-to-fine manner at 32×32×32 nm^3^, 16×16×16 nm^3^, and (native) 8×8×8 nm^3^ resolution in order to generate a base segmentation that was largely over-segmented. We then agglomerated the base segmentation, also using FFNs. We aggressively agglomerated segments despite introducing substantial numbers of erroneous mergers, as advances in proofreading methodology described elsewhere in this manuscript enabled efficient detection and correction of those mergers.

We evaluated the accuracy of the FFN segmentation of the hemibrain using expected run length (ERL) and merge rate metrics[17]. The base segmentation (i.e., the automated reconstruction prior to agglomeration) achieved an ERL of 163 microns with a merge rate of 0.25%. After (automated) agglomeration, run length increased to 585 microns but with a false merge rate of 27.56% (i.e., nearly thirty percent of of the ground truth path length was contained in automated segments, but these had, on average, at least one merge error). We also evaluated a subset of neurons in the volume, ~500 olfactory PN and KC cells chosen to roughly match the evaluation performed in [27] which yielded an ERL of 825 microns at a 15.92% merge rate.

### 2.3 Synapse Prediction

Accurate synapse identification is crucial since synapses both form a critical component of a connectome and are required for prioritizing and guiding the proofreading effort. Synapses in *Drosophila* are typically polyadic, with one pre-synaptic site (a T-bar) contacted by multiple receiving partners (often called PSDs, for post-synaptic densities) as shown in Fig 4a. Initial synapse prediction revealed that there are over 9 million T-bars and 60 million PSDs in the hemibrain. Manually validating each one, assuming a rate of 1000 connections annotated per trained person, per day, it would take more than 230 working years. Given this infeasibility, we developed machine learning approaches to predict synapses as detailed below. The results of this prediction are shown in Fig 4b, where the predicted synapse sites clearly delineate many of the fly brain regions. A major challenge from a machine learning perspective is the range of varying image statistics across the volume. In particular, model performance can quickly degrade in regions of the data set whose statistics are not well-captured by the training set[28]. To address this challenge, we took an iterative approach to synapse prediction, interleaving model re-training with manual proofreading, all based on the methods of [29]. Initial prediction, followed by proofreading, revealed a number of false positive predictions due to structures such as dense core vesicles which were not well-represented in the original training set. A second filtering network was trained on regions causing such false positives, and used to prune the original set of predictions. We denote this pruned output as the ‘initial’ set of synapse predictions.

**Figure 4:**
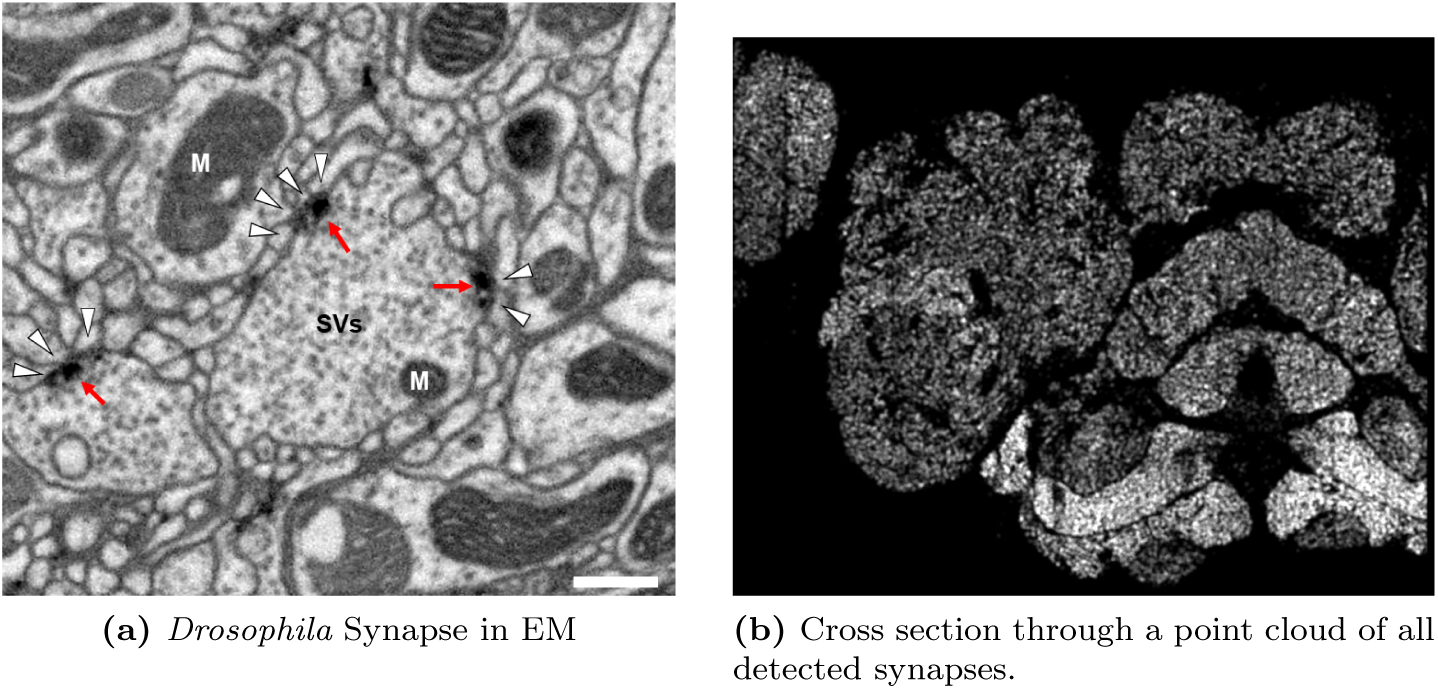
Well preserved membranes, darkly stained synapses, and smooth round neuronal processes are characteristics of the hemibrain sample. Panel (a) shows polyadic synapses, with a red arrow pointing to the pre-synaptic T-bar, and white triangles pointing to the post-synaptic densities. Mitochondria are labeled ‘M’, synaptic vesicles ‘SV’, and the scale bar is 0.5 *µ*m. Panel (b) shows a cross section through a point cloud of all detected synapses. This defines many of the compartments in the fly brain, much like an optical image obtained through nc82 antibody (an antibody against a component of T-bars) staining of synapses. This is used for generating the transformation from our sample to the standard *Drosophila* brain.

Based on this initial set, we began collecting human-annotated dense ground-truth cubes throughout the various brain regions of the hemibrain, to assess classifier performance variation by brain region. From these cubes, we determined that although many regions had acceptable precision, there were some regions in which recall was lower than desired. Consequently, a subset of cubes available at that time was used to train a new classifier focused on addressing recall in the problematic regions. This new classifier was used in an incremental (cascaded) fashion, primarily by adding additional predictions to existing initial set. This gave better performance than wholesale replacement, with the resulting predictions able to improve recall while largely maintaining precision.

As an independent check on synapse quality, we also trained a separate classifier proposed in [28], using an enhanced version of the ‘synful’ software package. Both synapse predictors also give a confidence value for each synapse, a measure of how firmly the classifier believes the found feature is a true synapse. We found that we were able to improve recall by taking the union of the two predictors most confident synapses, and similarly improve precision by removing synapses that were low confidence in both predictions. Figures 5a and 5b show the results, showing the precision and recall obtained in each brain region.

**Figure 5:**
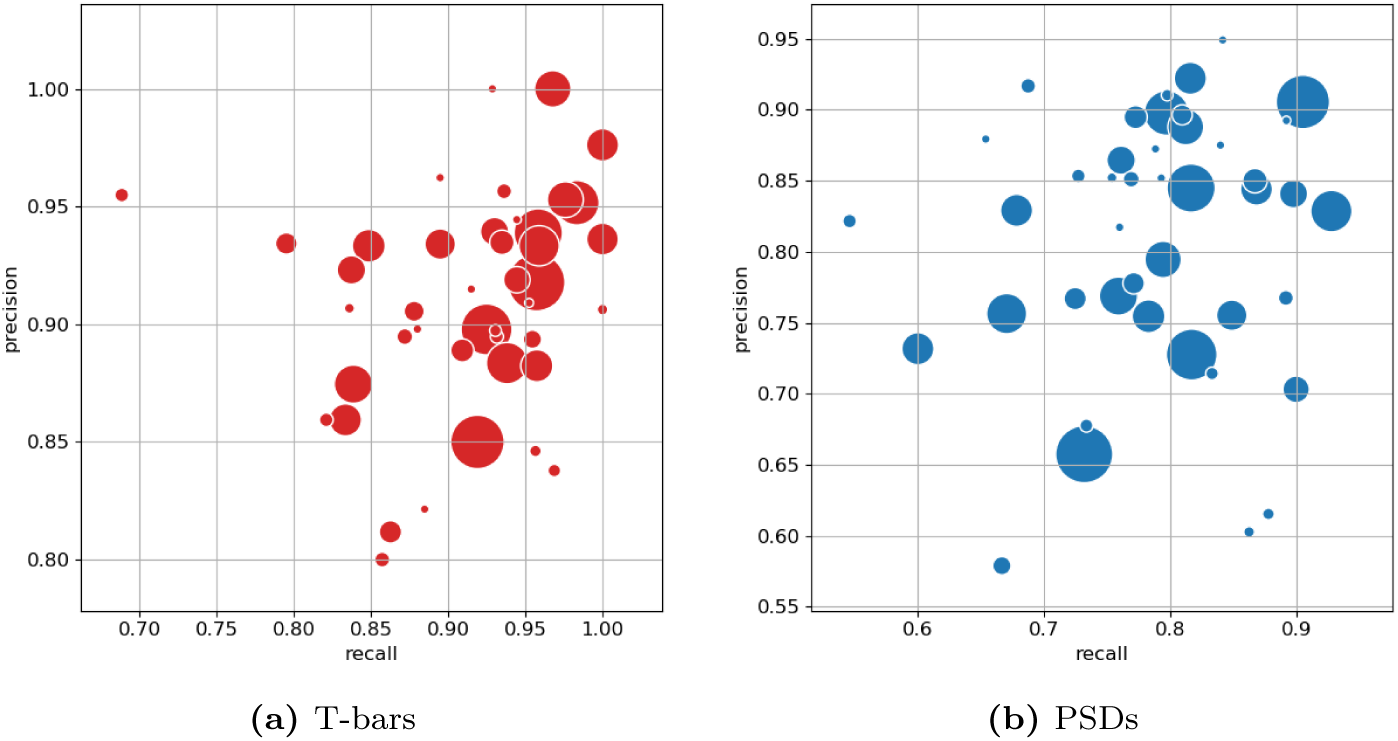
Precision and recall for synapse prediction. The plot on the left indicates precision and recall for T-bars; the plot on the right for synapses as a whole including PSDs identification. T-bar identification is better than PSD identification since the feature is both more distinct and typically occurs in larger neurites. Each dot is one brain region. The size of the dot is proportional to the volume of the region. Humans typically achieve 0.9 precision/recall on T-bars and 0.8 precision/recall on PSDs.

### 2.4 Proofreading

Since machine segmentation is not perfect, we made a concerted effort to fix the errors remaining at this stage by several passes of human proofreading. Segmentation errors can be roughly grouped into two classes - false merges, in which two separate neurons are mistakenly merged together, and false splits, in which a single neuron is mistakenly broken into several segments. Enabled by advances in visualization and semi-automated proofreading using our Neu3 tool[30], we first addressed large false mergers. A human examined each putative neuron and determined if it had an unusual morphology suggesting a merge might have occurred, a task still much easier for humans than machines. If judged to be a false merger, the operator identified discrete points that should be on separate neurons. The shape was then resegmented in real time allowing users to explore other potential corrections. Neurons with more complex problems were then scheduled to be re-checked, and the process repeated until few false mergers remained.

In the next phase, the largest remaining pieces were merged into neuron shapes using a combination of machine-suggested edits[31] and manual intuition, until the main shape of each neuron emerged. This requires relatively few proofreading decisions and has the advantage of producing an almost complete neuron catalog early in the process. As discussed below, in the section on validation, emerging shapes were compared against genetic/optical image libraries (where available) and against other neurons of the same putative type, to guard against large missing or superfluous branches. These procedures (which focused on higher-level proofreading) produced a reasonably accurate library of the main branches of each neuron, and a connectome of the stronger neuronal pathways. At this point there was still considerable variations among the brain regions, with more completeness achieved in areas where the initial segmentation performed better.

Finally, to achieve highest reconstruction completeness possible in the time allotted, and to enable confidence in weaker neuronal pathways, proofreaders connected remaining isolated fragments (segments) to already constructed neurons, using NeuTu[32] and Neu3[30]. The fragments that would result in largest connectivity changes were considered first, exploiting automatic guesses through focused proofreading where possible. Since proofreading every small segment is still prohibitive, we tried to ensure a basic level of completeness throughout the brain with special focus in regions of particular biological interest such as the central complex and mushroom body.

### 2.5 Defining brain regions

In a parallel effort to proofreading, the sample was annotated with discrete brain regions, as shown in Fig. 6. This process used the criteria of synapse density, glial boundaries, neural tracts, and detailed neuron wiring, similar in spirit to the methods of [1]. Synapse density was computed as a point cloud of synapse locations, yielding results similar to nc82 staining but more precise, as shown in Fig 4b above. Neural tracks were identified manually. Machine learning was used to identify glia with EM level precision.

**Figure 6:**
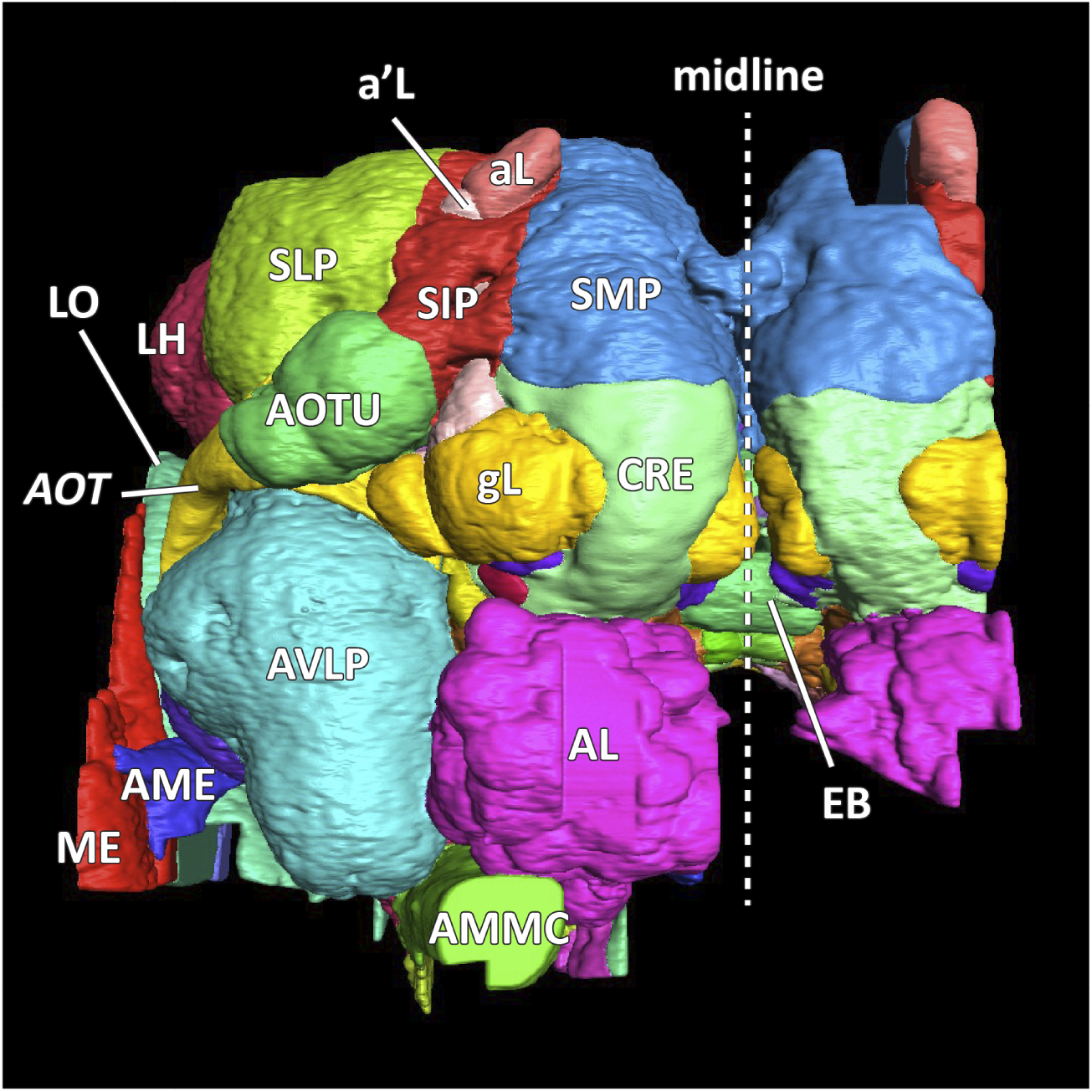
A frontal (anterior) view of the reconstructed brain regions of the hemibrain dataset.

In some cases, we also defined substructures, such as layers and slices, within the central complex and mushroom bodies, although their boundaries are often imprecise because of the lack of clear landmarks. They were defined with reference to traditional genetic and optical methods, using known cell types as references. To avoid double counting, substructures were defined to be mutually exclusive. This means that (as is also true in the optic lobe) that a neuron with arbors predominantly in one substructure may also have some synapses and smaller arbors in adjacent substructures.

### 2.6 Cell Type Classification

Defining cell types for groups of similar neurons is a time-honored mechanism to attempt to understand the anatomical and functional properties of a circuit. Presumably, neurons of the same type execute similar circuit roles. However, the definition of what is a distinct cell type and the exact delineation between one cell type and another is inherently vague and represents a classic taxonomic challenge, pitting ‘lumpers’ vs ‘splitters’. Despite our best efforts, we recognize that our typing is not exact, and expect ongoing revisions to cell type classification.

One common method of cell type classification, used in flies, exploits the GAL4 system to highlight the morphology of neurons with similar gene expression[33]. Since these genetic lines are imaged using fluorescence and confocal microscopy, we refer to them as ‘light lines’. Where they exist and are sufficiently sparse, light lines provide a key method for identifying types by grouping morphologically similar neurons together. However, there are several limitations. There are no guarantees of coverage, and it is sometimes difficult to distinguish between neurons of very similar morphology.

We enhanced the classic view of morphologically distinct cell types by defining distinct cell types (or sub-cell type groupings, if one prefers) based on connectivity as well as morphology. This connectivity-based clustering often serves a clear arbiter of cell type distinction, even when genetic markers have not yet been found, or when the morphology of different types is quite similar, sometimes sufficiently similar to be indistinguishable in optical images. For example, the two PEN neurons below have very similar morphology but quite distinct inputs, as shown in Fig 7 (PEN1 and PEN2 neurons, in fact, have been shown to have different functional activity in [34]).

**Figure 7:**
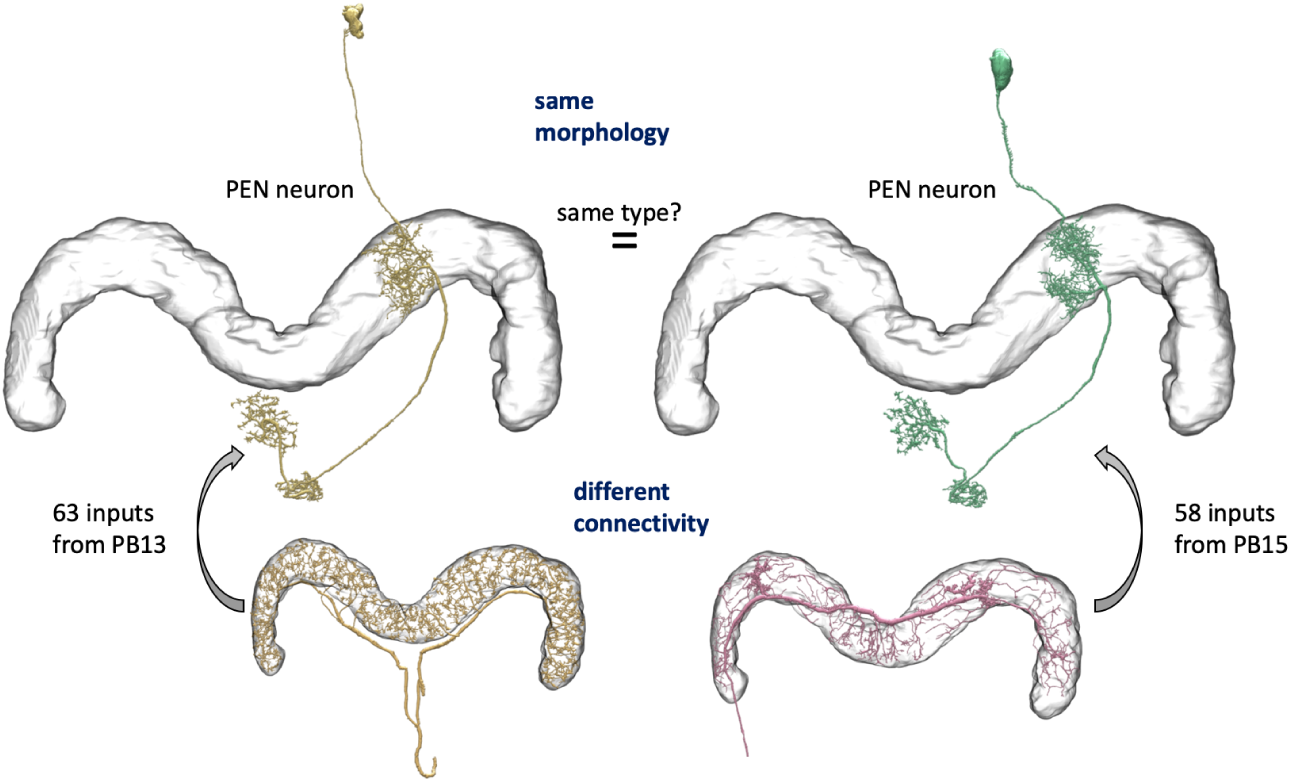
An example of two neurons with very similar morphology but different connectivity.

### 2.7 Methodology for assigning cell types and nomenclature

Assigning names and types to the more than 20,000 reconstructed cells was a difficult and contentious undertaking. Most of the neurons have no previously annotated type. Adding to the complexity, prior work focused on morphological similarities and differences, but here we have, for the first time, connectivity information to use in typing as well.

Many cell types are already described in the literature, but the existing names can be both inconsistent and ambiguous. The same cell type is often given differing names in different publications, and conversely, the same name, such as PN for projection neuron, is used for many different cell types. Nonetheless, for cell types already named in the literature (famous cell types), we have tried to use an existing name. We apologize in advance for any offense given by our selection of names.

Overall, we defined a ‘type’ of neurons as a single or a group of cells that have very similar cell body location, morphology, and synaptic connectivity patterns, and employed a four-step process to classify them. We found 18,478 neuronal cell bodies in the hemibrain data, among them 15,084 are on the right side of the central brain, whereas the rest are in the right optic lobe or on the left side of the central brain near the midline.

The first step was to classify all cells by their lineage. We first grouped neurons according to their bundle of cell body fibers (CBFs). Neuronal cell bodies are located in the cell body rind around the periphery of the brain, and each neuron projects a single CBF towards synaptic neuropils. In the central brain, cell bodies of clonally related neurons deriving from a single stem cell tend to form clusters, which sends one or several bundles of CBFs. We carefully examined the trajectory and origins of CBFs of the 15,084 neurons on the right central brain and identified 192 distinct CBF bundles. Among them, 154 matched the CBF bundles of 102 known clonal units[35][36]. The rest are minor populations and most likely of embryonic origin. Each of the 192 CBF bundles was given an ID according to the location of the cell body cluster (split into eight sectors of the brain surface with the combination of Anterior/Posterior, Ventral/Dorsal, and Medial/Lateral) and a number within the sector given according to the size of cell population. Thus, a CBF group might be named ADM01, meaning a group with the largest number of neurons in the Anterior Dorsal Medial sector of the brain surface.

Different stem cells sometimes give rise to neurons with very similar morphology. We classified these as different types because of their distinct developmental origin. Thus, the second step of neuron typing was to cluster neurons within each CBF group. This process consisted of three further steps. First, we subjected all the neurons of a particular CBF group to morphology-based clustering using NBLAST[37]. Next, the resulting cell grouping was used as a template to initiate iterative connectivity-based clustering. This process sometimes re-grouped neurons with different morphology, if they shared similar connectivity patterns. Finally, the clustering results were subject to extensive manual review and visual inspection, and compared using NBLAST dendrograms to ensure that each cell type consisted of morphologically similar neurons. This review allowed us to both confirm cell type identity and helped ensure reconstruction accuracy.

In the hemibrain, using the defined brain regions and reference to known light lines, biologists were able to assign a cell type to many cells. Where possible, we matched previously defined cell types with those labeled in light data using a combination of NBLAST[37], Neuprint (described below), and human intuition to find the matching cell types, especially in well explored regions such as the mushroom body (MB) and central complex (CX), where abundant cell type information was already available and where we are more confident in our anatomical expertise. Even though most of the cell types in MB and CX were already described, we still found new cell types, and tried to name them using existing schemes for these regions. We further refined these morphological groupings with connectivity information when relevant.

However, outside the heavily studied regions the fly’s circuits are largely composed of cells of unknown type. In this case putative type names were derived from the CBF group, morphological type, and connectivity type. Morphological types were represented by the CBF group name followed by an underscore and 1-3 lowercase letters. If neurons of near-identical morphology could be further subdivided into different connectivity types, they were suffixed with an underscore and an uppercase letter. Finally, a suffix ‘ pct’, for putative cell type, was added. Thus, a full putative type name might be ‘ADM01a pct’ if all the neurons of this type shared similar connectivity patterns, or ‘ADM01b A pct’ if there are different connectivity types within neurons with similar morphology. The resulting names are not pretty, but the process is systematic and scalable. The assignment of type names to neurons is still ongoing, and we expect the names of putative cell types will be refined by the research community, including simpler and easier to pronounce names, as new information emerges. What will **not** change are the unique body ID numbers given in the database that refer to a particular (traced) cell in this particular image dataset. We strongly advise that such IDs be included in any publications based on our data to avoid confusion as cell type names (and possibly instance names) evolve.

### 2.8 Results of cell typing

Using the above semi-automated procedures, we identified 4,768 cell types for 20,607 neurons. Over a thousand of these are types with only a single instance (i.e only a single neuron), although presumably, for a whole brain reconstruction, most of these types would have a matching instance in the other side of the brain. Fig. 8 below shows the number of distinct neuron types found in different brain regions. Fig. 9 shows the distribution of the number of neurons in each cell type.

**Figure 8:**
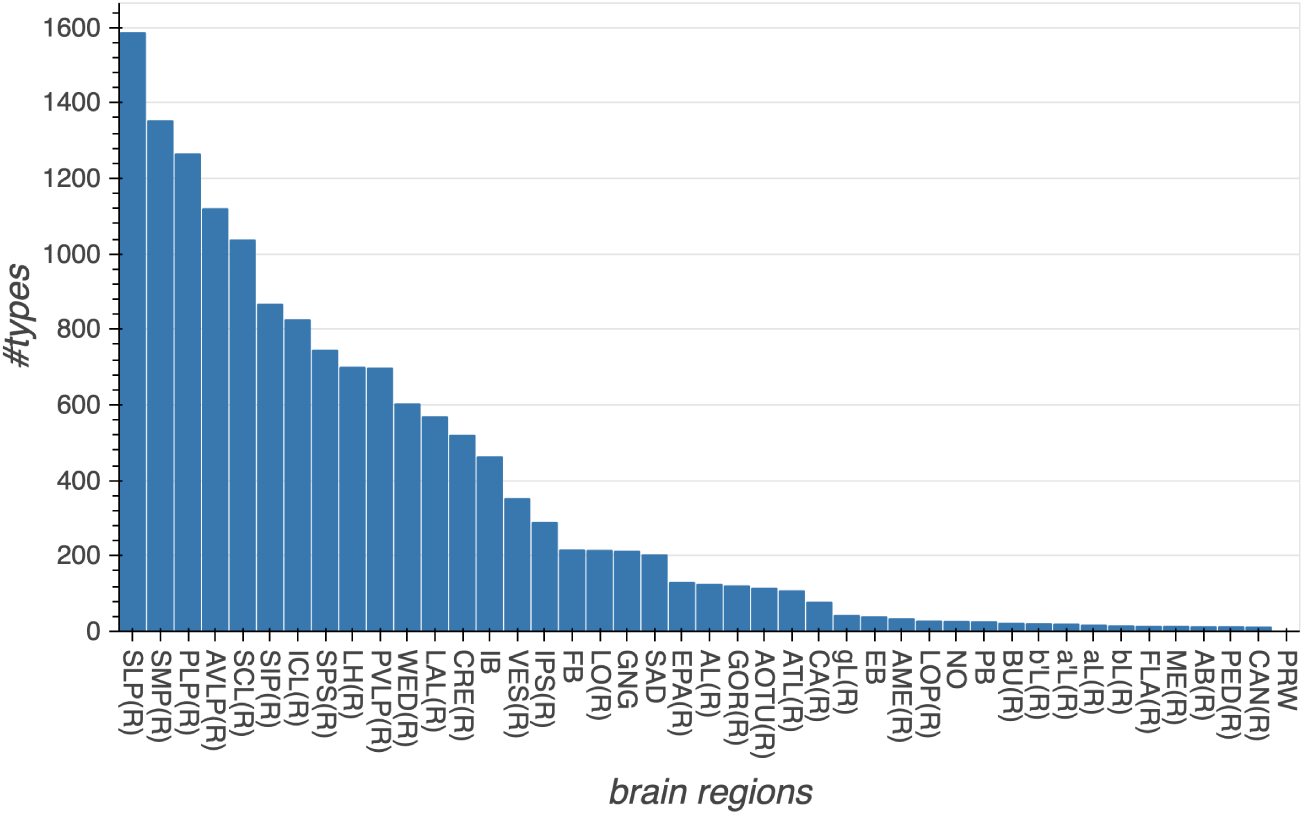
The number of cell types in each major brain region.

**Figure 9:**
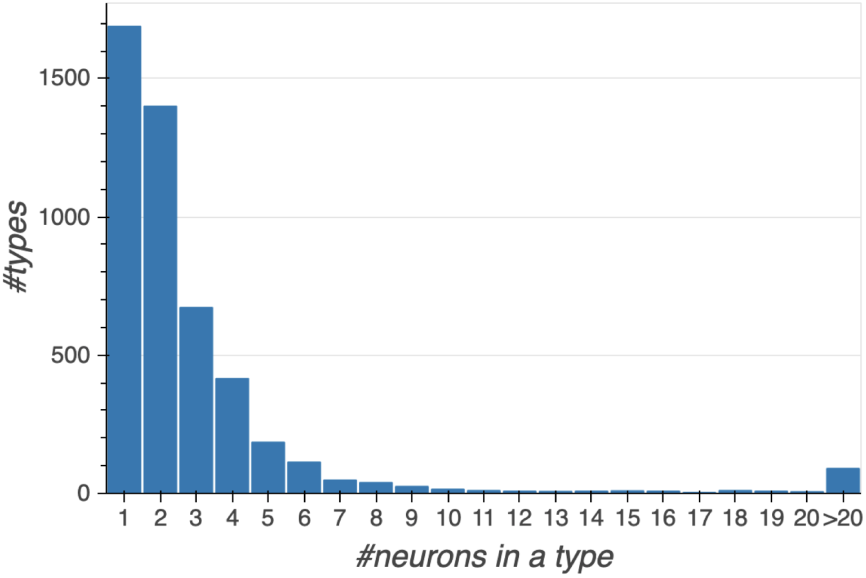
Histogram showing the number of cell types with a given number of constituent cells.

### 2.9 Assessing Morphologies and Cell Types

Verifying correctness and completeness is a challenging problem since there is no existing full brain connectome against which our data might be compared. We devised a number of tests to check the main features: Are the morphologies correct? Are the regions and cell types correctly defined? Are the synaptic connection counts representative?

Assessing completeness is much easier than assessing correctness. Since the reconstruction is dense, we believe the census of cells, types, and regions is essentially complete. The main arbors of every cell within the volume are reconstructed, and almost every cell is assigned to at least a putative cell type. Similarly, since the identified brain regions nearly tile the brain, these are complete as well.

For checking morphologies, we searched for major missing or erroneous branches using a number of heuristics. Each neuron was reviewed by multiple proofreaders. The morphology of each neuron was compared to light data whenever it was available. When more than one cell of a given type was available, a human compared the two, which helped us find missing or extra branches, and also served as a double check on the cell type assignment. In addition, since the reconstruction is dense, all sufficiently large orphan neurites were examined manually until they were determined to form part of a neuron, or they left the volume. To help validate the assigned cell types, proofreaders did pairwise checks of every neuron of similarly scored types.

For subregions where previous dense proofreading was available (such as the alpha lobes of the mushroom body) we compared the two connectomes. We were also helped by research groups using both sparse tracing in the full fly brain TEM dataset[6], and our hemibrain connectome. They were happy to inform us of any inconsistencies. There are limits to this comparison, as the two samples were different ages and raised under different conditions, then prepared and imaged by different techniques, but this likely would have revealed any gross errors. Finally, we generated a probabilistic connectome based on a different segmentation, and systematically visited regions where the two differed.

### 2.10 Assessing Synapse Accuracy

As discussed in the section on finding synapses, we evaluated both precision (are the found synapses real?) and recall (did we find all the synapse that exist?) on sample cubes in each brain region. We also double checked by comparison with a different synapse detection algorithm.

As a final check, we also evaluated the end-to-end correctness of given connections between neurons for different cell types and across brain regions. This means we checked that the pre and post synaptic annotations were correct, and also whether they were assigned to the correct neuron. For the worst few examples, we manually refined synapse predictions. Fig 10 shows the results: we were able to obtain high precision for most cell types with a relatively small number of lower precision outliers. We also evaluated single-connection pathways across each brain region. In the fly, important connections typically have many synapses between them. However, the presence of connections represented by low numbers of synapses is well known, although their biological importance is unclear. Regardless, we wanted to ensure that even single connection pathways were mostly correct. Our analysis suggests that this is indeed the case over 76% of the time, a number that does not seem to vary greatly between brain regions.

**Figure 10:**
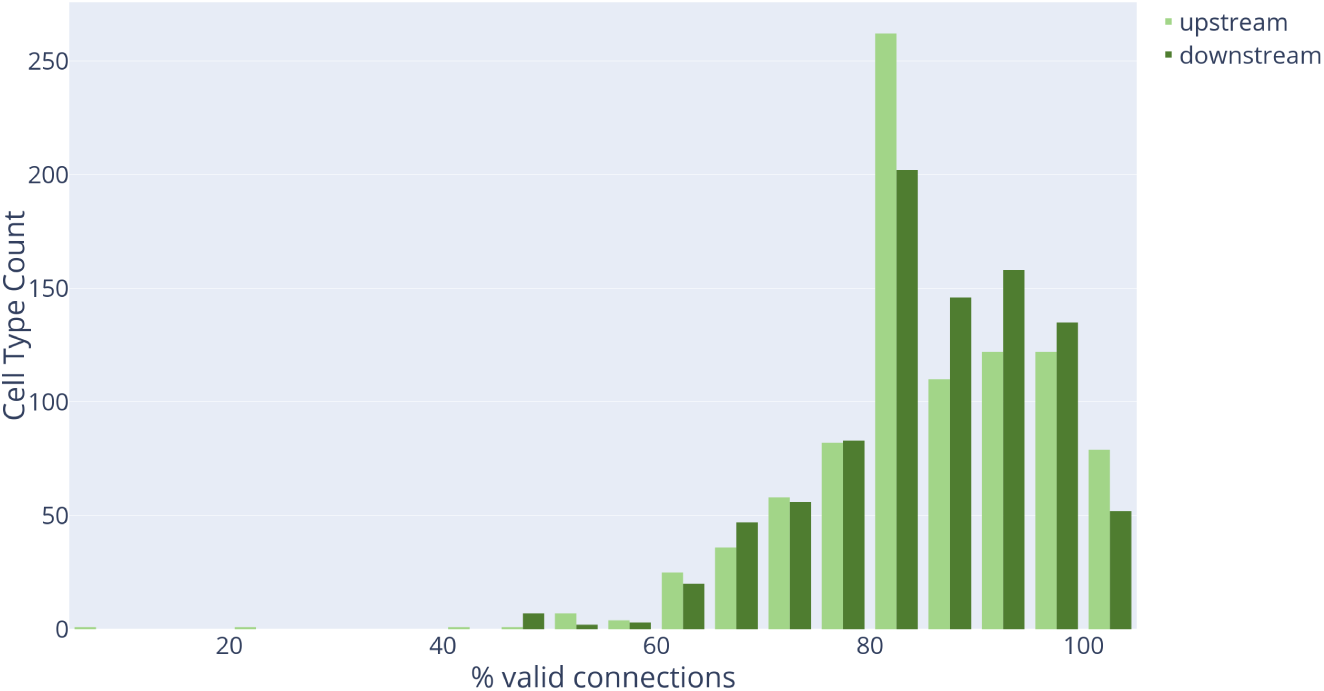
Connection precision of upstream and downstream partners for 1000 cell types.

### 2.11 Assessing connection completeness

A synapse in the fly brain consists of a pre-synaptic density (T-bar) and typically several post-synaptic partners (PSDs). The T-bars are contained in larger neurites, and most (>90%) of the T-bars in our dataset are contained in identified neurons. The post-synaptic densities are typically in smaller neurites, and it is these that are difficult for machine (or human) to connect with certainty.

With current technology, tracing all fine branches in our EM images is impractical, so we sample among them (at completeness levels typically ranging from 20% to 85%) and trace as many as practical in the allotted time. The goal is to provide synapse counts that are representative, since completeness is out of reach. Provided the missing PSDs are independent (which we try to verify), then the overall circuit emerges even if a substantial fraction of the connections are missing. If a connection has a strength of 10, for example, then it will be found in the final circuit with more than 99.9% probability, provided at least half the individual synapses are traced.

If unconnected small twigs are the main source of uncertainty in our data (as we believe to be the case), then as proofreading proceeds existing connections should only get stronger. Of course corrections resulting in lesser connection strength, such as fixing a false connection or removing an incorrect synapse, are also possible, but are considerably less likely. To see if our proofreading process is working as we expect, we take a region that has been read to a lower percentage completion and then spend the manual effort to reach a higher percentage, and then compare the two circuits. (A versioned database such as DVID is enormously helpful here.) If our efforts are successful, ideally what we see is that almost all connections that change get stronger, very few connections get weaker, and no new strong connections appear (since all strong connections should already be present even in low coverage proofreading). If this is the behavior we find, we can be reasonably sure that the circuits found are representative for all strong connections.

Fig. 11 below shows such an analysis. The results support our view that the circuits we report reflect what would be observed if we extrapolated to assignment of all pre- and post-synaptic elements.

**Figure 11:**
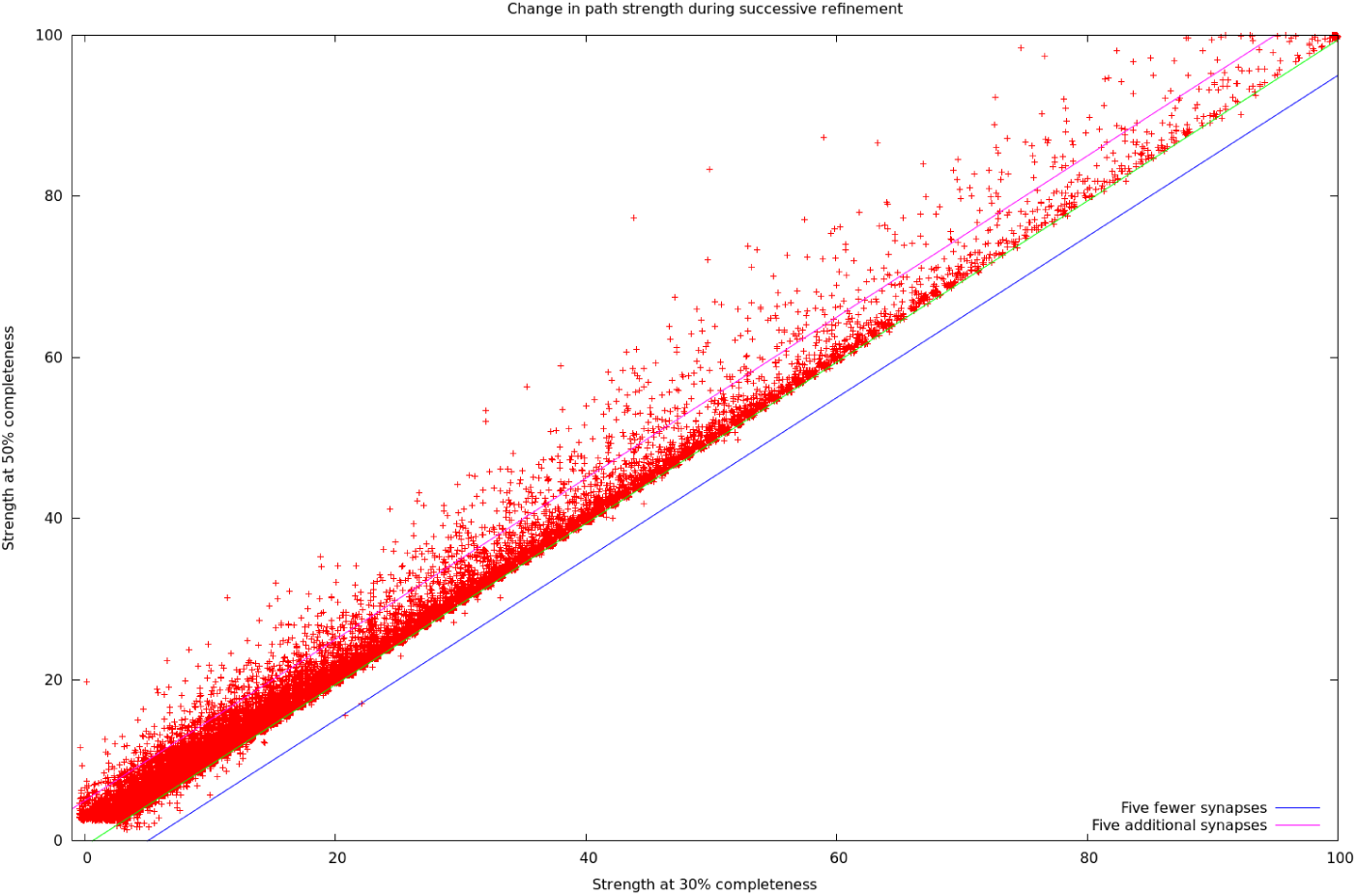
Difference between connection strengths in the Ellipsoid Body as proof-reading completeness increased. Roughly 40,000 paths are shown. Almost all points are above the line Y=X, showing that almost all paths increased in strength, with very few decreasing. In particular, no path decreased in strength by more than 5 synapses. Only two new strong (strength > 10) paths were found that were not present in the original. This should be rarer at higher levels of proofreading since neuron fragments (orphans) are added in order of decreasing size (see text).

### 2.12 Interpreting the connection counts

Given the complexity of the reconstruction process, and the many different errors that could occur, how confident should the user be that the returned synapse counts are valid? This section gives a quick guide in the absence of detailed investigation. The number of synapses we return is the number we found. The true number could range from slightly less, largely due to false synapse predictions, to considerably more, in the regions with low percentage reconstructed. For connections known to be in a specific brain region, the reciprocal of the completion percentage (as shown in Table 1) gives a reasonable estimate of undercount.

If we return a count of 0 (the neurons are not connected), there are two cases. If the neurons do not share any brain regions, then the lack of connections is real. If they do share one or more brain regions, then a count of 0 is suspect. It is possible that there might be a weak connection (count 1-2) and less likely there is a medium strength connection (3-9 synapses). Strong connections can be confidently ruled out, minus the small chance of a mis- or unassigned branch with many synapses.

If we report a weak connection (1-2 synapses), then the true strength might range from 0 (the connection does not exist) up through a weak connection (3-9 synapses). If your model or analysis relies on the strength of these weak connections, its a good idea to manually check our reconstruction. If your analysis does not depend on knowledge of weak connections, we recommend ignoring connections based on 3 or fewer synapses.

If we report a medium strength connection (3-9 synapses) then the connection is real. The true strength could range from weak to the lower end of a strong connection.

If we report a strong connection (10 or more synapses), the connection not only exists, but is strong. It may well be considerably stronger than we report.

## 3 Data Representation

The representation of connectomics data is a significant problem for all connectomics efforts. The raw image data on which our connectome is based is larger than 20 TB, and takes 2 full days to download even at a rate of 1 gigabit/second. Looking forward, this problem will only get worse. Recent similar projects are generating petabytes worth of data[38], and a mouse brain of 500 mm^3^, at a typical FIB-SEM resolution of 8nm isotropic, would require almost 1000 PB.

In contrast, most users of connectivity information want a far smaller amount of much more specific information. For example, a common query is ‘what neurons are downstream (or upstream) of a given target neuron?’. This question can be expressed in a few tens of characters, and the desired answer, the top few partners, fits on a single page of text.

Managing this wide range of data, from the raw gray-scale through the connectivity graph, requires a variety of technologies. An overview of the data representations we used to address these needs is shown in Fig. 12: This organization offers several advantages. In most cases, instead of transferring files, the user submits queries for the portion of data desired. If the user needs only a subset of the data (as almost all users do) then they need not cope with the full size of the data set. Different versions of the data can be managed efficiently behind the scenes with a versioned database such as DVID[39] that keeps track of changes and can deliver data corresponding to any previous version. The use of existing software infrastructure, such as Google buckets or the graph package neo4j, which are already optimized for large data, helps with both performance and ease of development. The advanced user is not limited to these interfaces – for those who wish to validate or extend our results, we have provided procedures whereby the user can make personal copies of each representation, including the grayscale, the DVID data storage, and our editing and proofreading software. This allows other researchers to establish an entirely independent version of all we have done, completely under their control. Contact the authors for the details of how to copy all the underlying data and software.

**Figure 12:**
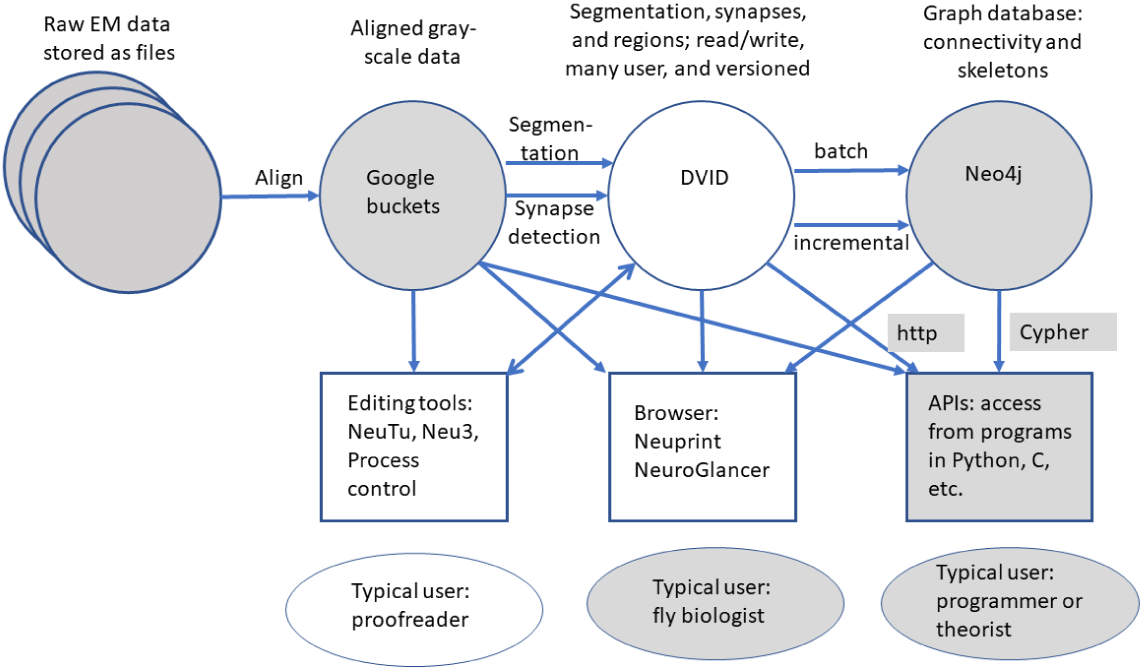
Overview of data representations of our reconstruction. Circles are stored data representations, rectangles are application programs, ellipses represent users, and arrows indicate the direction of data flow labelled with transformation and/or format. Filled areas represent existing technologies and techniques; open areas were developed for the express purpose of EM reconstruction of large circuits.

### 3.1 What are the data types?

Grayscale data correspond to traditional electron microscope images. This is written only once, after alignment, but often read, as it is required for segmentation, synapse finding, and proofreading. We store the grayscale data, 8 bits per voxel, in Google buckets, which makes access from geographically distributed sites easier.

Segmentation, synapses, and regions annotate and give biological meaning to the grayscale. For segmentation, we assign a 64 bit neuron ID to each voxel. Despite the larger size per voxel (64 vs 8 bits) compared to the grayscale, the storage required is much smaller (by a factor of more than 20) since segmentation compresses well. Although the voxel level segmentation is not needed for connectivity queries, it may be useful for tasks such as computing areas and cross-sections at the full resolution available, or calculating the distance between a feature and the boundary.

Synapses are stored as point annotations - one point for a pre-synaptic T-bar, and one point for each of its post-synaptis densities (or PSDs). The segmentation can then be consulted to find the identity of the neurons containing the synapses.

The compartment map of the brain is stored as a volume specified at a lower resolution, typically a 32×32×32 voxel grid. At 8nm voxels, this gives a 256 nm resolution for brain regions, comparable to the resolution of confocal laser scanning microscopy.

Unlike the gray scale data, segmentation, synapses, and regions are all modified during proofreading. This requires a representation that must cope with many users modifying the data simultaneously, log all changes, and be versioned. We use DVID[39], developed internally, to meet these requirements.

Neuron skeletons are computed from the segmentation[40], and not entered or edited directly. A skeleton representation describes each neuron with (branching) centerlines and diameters, typically in the SWC format popularized by the simulator *Neuron*[41]. These are necessarily approximations, since it normally not possible (for example) to match both the cross sectional area and the surface area of each point along a neurite with such a representation. But SWC skeletons are a good representation for human viewing, adequate for automatic morphology classification, and serve as input to neural simulations such as Neuron. SWC files are also well accepted as an interchange format, used by projects such as NeuroMorpho[42] and FlyBrain[43].

The connectivity graph is also derived from the data and is yet more abstract, describing only the identity of neurons and a summary of how they connect - for example, Neuron ID1 connects to neuron ID2 through a certain number of synapses. In our case it also retains the brain region information and the location of each synapse. Such a connectivity graph is both smaller and faster than the geometric data, but sufficient for most queries devised by biologists, such as finding the upstream or downstream partners of a neuron. A simple connectivity graph is often desired by theorists, particularly within brain regions, or when considering neural circuits where each neuron can be represented as a single node.

A final, even more abstract form is the adjacency matrix: This compresses the connectivity between each pair of neurons to a single number. Even this most economical form requires careful treatment in connectomics. As our brain sample contains more than 25K traced neurons as well as many unconnected fragments, the adjacency matrix has more than a billion entries (most of which are zero). Sparse matrix techniques, which report only the non-zero coefficients, are necessary for practical use of such matrices.

## 4 Accessing the data

For this project we provide access to the data through a combination of a software interface[2] and a server (https://neuprint.janelia.org). These allow access to not just the current project data but to two previous connectomics efforts as well (a 7-column optic lobe reconstruction[44] and the alpha lobe of the mushroom body[4]). Data are available in the form of gray-scale, pixel-level segmentation, skeletons, and a graph representation.

The most straightforward way to access the hemibrain data is through the Neuprint[2] interactive browser. This is a web-based application that is intended to be usable by biologists with minimal or no training. It allows the selection of neurons by name, type, or brain region, displays neurons, their partners, and the synapses between them in a variety of forms, and provides many of the graphs and summary statistics that users commonly want.

Neuprint also supports queries from languages such as Python[45] and R, as used by the neuroanatomy tool NatVerse[46]. Various formats are supported, including SWC format for the skeletons. In particular, the graph data can be queried through an existing graph query language, Cypher[47], as seen in the example below. The schema for the graph data is shown in Fig. 13.

> MATCH (n:Neuron) - [c:ConnectsTo] -> (t:Neuron) WHERE t.type = ‘MBON18’
>
> RETURN n.type, n.bodyId, c.weight ORDER BY c.weight DESCENDING

**Figure 13:**
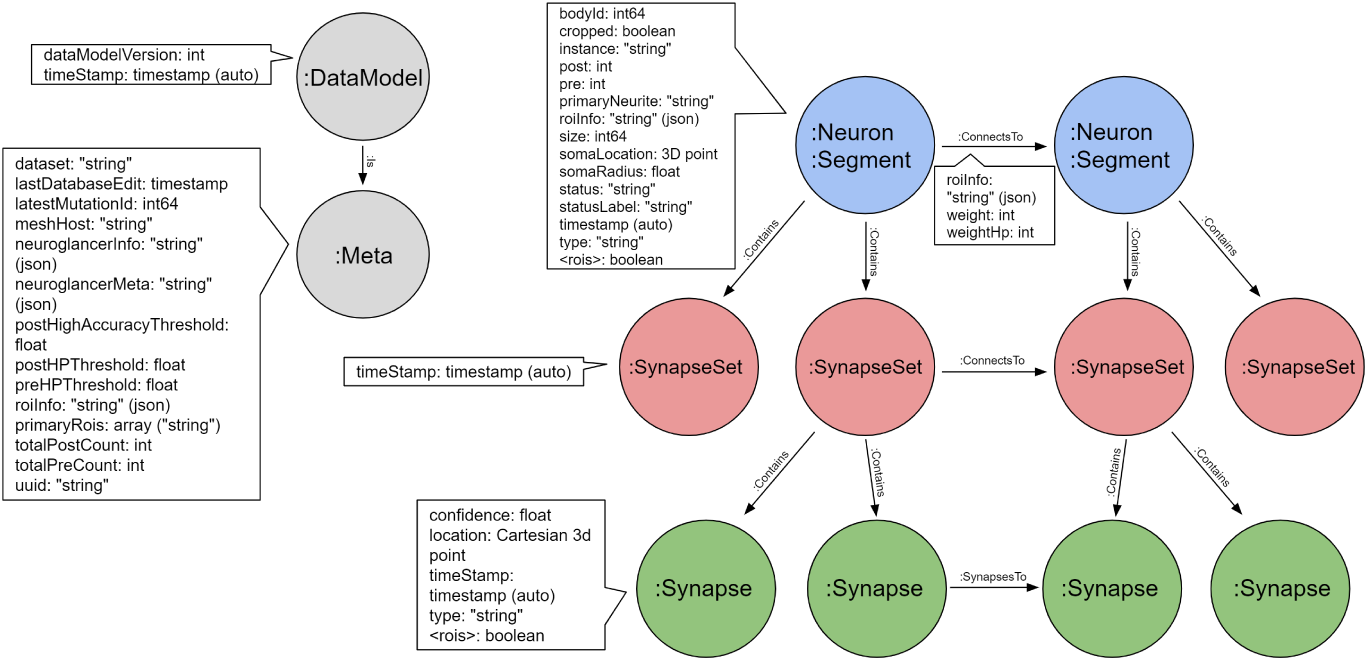
Schema for the neo4j graph model of the hemibrain. Each neuron contains 0 or more SynapseSets, each of which contains one or more synapses. All the synapses in a SynapseSet connect the same two neurons. If the details of the synapses are not needed, the neuron to neuron weight can be obtained as a property on the ConnectsTo relation, as can the distribution of this weight across brain regions (the roiInfo).

This query looks for all neurons that are pre-synaptic to any neuron of type ‘MBON18’. For each such neuron it returns the types and internal identities of the pre-synaptic neuron, and the count of synapses between them. The whole list is ordered in order of decreasing count. This is just an illustration; this particular query is quite common and is supported in Neuprint without the need for any programming language.

Adjacency data, if needed, can be derived from the graph representation. We provide a small demonstration program that queries the API and generates such matrices, either with or without the brain regions. The two matrices themselves are available in gzipped Python format. For more information on accessing data and other hemibrain updates, please see https://www.janelia.org/project-teams/flyem/hemibrain.

### 4.1 Matching EM and light data

We registered the hemibrain EM data to the JRC2018 *Drosophila* template brain[48] using an automatic registration algorithm followed by manual correction. We began by using the automated T-bar predictions (described in section 2.3) to generate a T-bar density volume rendered at a resolution comparable to light microscopy images. This hemibrain synapse density volume was automatically registered to the template brain using ANTs[49], producing both a forward and inverse transform. The resulting registration was manually fine-tuned using BigWarp[50]. The total transform is the composition of the ANTs and BigWarp transformations.

Given a particular neuron of interest, researchers can use these resources to identify GAL4 lines labeling that neuron. First the representation of the neuron must be spatially transformed into the template space that GAL4 driver line images have been registered to. A mask based approach[51] enables searching GAL4 driver line image databases for particular neurons. Skeletonizing hemi-brain neurons can enable querying GAL4 neuronal skeleton databases using NBLAST[37].

### 4.2 Longer term storage of data, and archival references

Historically, archival biology data have been expressed as files often included with supplementary data. However, for connectivity data this has two main problems. First, the data are large, and hard to store. Journals, for example, typically limit supplemental data to a few 10s of megabytes. The data here is about 6 orders of magnitude larger. Second, connectome data is not static, during proofreading and even after initial publication. As proofreading proceeds, the data improves in completeness and quality. The question then is how to refer to the data as it existed at some point in time, required for reproducibility of scientific results. If represented as files, this would require many copies, check-pointed at various times - the as submitted version, the as published version, the current best version, and so on.

We resolve this, at least for now, by hosting the data ourselves and making it available through query mechanisms. Underlying our connectome data is a versioned database (DVID) so it is technically possible to access every version of the data as it is revised. However, as it requires effort to host and format this data for the Neuprint browser and API, only selected versions (called named versions) are available from the website by default. The initial version is hemibrain:v1.0. Although this is only version currently, when reproducibility is required, such as referencing the data in a paper, it is still best to refer explicitly to the milestone versions by name (such as hemibrain:v1.0), as we expect a new milestone version every few months, at least at first. We will supply a DOI for each of these versions, and each is archived, can be viewed and queried through the web browser and APIs at any time, and will not change.

The goal of multiple versions is that later versions should be of higher quality. Towards this end we have implemented several systems for reporting errors so we can fix them. Users can add annotations in NeuroGlancer[52], the application used in conjunction with Neuprint to view image data, where they believe there are errors. To make this easier, we provide a video explaining this process. We will review these annotations and fix those that we agree are problems. Users can also contact us via email about problems they find.

Archival storage is an issue since, unlike genetic data, there is no institutional repository for connectomics data and the data are too large for journals to archive. We pledge to keep this data available for at least the next 10 years.

## 5 Analysis

Of necessity, most previous analyses have concentrated on particular circuits, cell types, or brain regions. For example, a classic paper about motifs[53] sampled the connections between one cell type (layer 5 pyramidal neurons) in one brain region (rat visual cortex), and found a number of non-random features, such as over-represented reciprocal connections and a log-normal strength distribution. However, it has never been clear which of these observations generalize to other cell types, other brain regions, and the brain as a whole. We are now in a position to make much stronger statements, ranging over all brain regions and cell types.

In addition, many analyses are best performed (or can only be performed) on dense connectomes. Type-wide observations (all cells of a certain type have property X) depend on a complete census of that cell type, and depending on the observation, a complete census of upstream and downstream partners as well. Some analyses, such as null observations about motifs (where certain motifs are not found in the fly brain) can only be done on dense connectomes.

### 5.1 Compartment statistics

One analysis enabled by a dense whole-brain reconstruction involves the comparison of the circuit architectures of different brain areas within a single individual. The compartments vary considerably. Table 2 shows the connectivity statistics of compartments that are completely contained within the volume, have at least 100 neurons, and have the largest or smallest value of various statistics. Across regions, the number of neurons varies by a factor of 74, the average number of partners of each neuron by a factor of 36, the network diameter by a factor of 4, the average strength of connection between partner neurons by a factor of 5, and the fraction of reciprocal connections by a factor of 5. The average graph distance between neurons is more conserved differing by a factor of only 2.

**Table 2:** Regions with minimum or maximum characteristics, picked from those regions completely within the reconstructed volume and containing at least 100 neurons. Yellow indicates a minimum value; green a maximal value. N is the number of neurons in the region, L is the number of connections between those neurons, <k> is the average number of partners (in the region), D is the network diameter, <str> is the average connection strength, broken up into non-reciprocal and reciprocal. fracR is the fraction of connections that are reciprocal, and AvgDist is the average number of hops (one hop corresponds to a direct synaptic connection) between any two neurons in the compartment. Network diameter is computed on the un-directed graph; all other metrics use the directed graph.

| Name | N | L | <k> | D | <str> | <non-r> | <r> | fracR | AvgDist |
| --- | --- | --- | --- | --- | --- | --- | --- | --- | --- |
| MB(R) | 3430 | 573664 | 167.249 | 8 | 3.270 | 3.076 | 3.383 | 0.632 | 2.189 |
| bL(R) | 1159 | 108134 | 93.299 | 8 | 2.019 | 1.855 | 2.122 | 0.613 | 2.090 |
| EB | 518 | 58793 | 113.500 | 4 | 10.053 | 4.627 | 12.172 | 0.719 | 1.756 |
| PLP(R) | 6689 | 224985 | 33.635 | 16 | 2.692 | 2.403 | 3.685 | 0.226 | 3.170 |
| SNP(R) | 9121 | 775474 | 85.021 | 13 | 2.987 | 2.515 | 4.492 | 0.239 | 2.748 |
| RUB(L) | 123 | 576 | 4.683 | 6 | 7.682 | 2.852 | 20.686 | 0.271 | 2.743 |
| EPA(R) | 1468 | 18199 | 12.397 | 13 | 2.171 | 2.098 | 2.644 | 0.134 | 3.496 |

## 6 Conclusions and future work

In this work we have achieved a dream of anatomists that is more than a century old. For at least the central brain of at least one animal with a complex brain and sophisticated behavior, we have a complete census of all the neurons and all the cell types that compose the brain, a definitive atlas of the regions in which they reside, and a graph representing how they are connected.

To achieve this, we have made improvements to every stage of the reconstruction process. Better means of sample preparation, imaging, alignment, segmentation, synapse finding, and proofreading are all summarized in this work and will form the basis of yet larger and faster reconstructions.

We have provided the data for all the circuits of the central brain, at least as defined by nerve cells and chemical synapses. This includes not only circuits of regions that are already the subject of extensive study, but also a trove of circuits whose structure and function are yet unknown.

Finally, we have provided a public resource that should be a huge help to all who study fly neural circuits. Finding upstream and downstream partners, a task that typically took months of finicky experiments, is now replaced by a lookup on a publically available web site. Detailed circuits, which used to require considerable patience, expertise, and expertise to acquire, are now available for the cost of an internet query.

Many of the extensions of this work are obvious and already underway. Not all regions of the hemibrain have been read to the highest accuracy possible, as we have concentrated first on the regions overlapping with other projects, such as the central complex and the mushroom body. We will continue to update other sections of the brain, and distributed circuits such as clock neurons that are not confined to one region, but spread throughout the brain.

Next, reconstructing a male fly is critical since the circuits of the two sexes are known to differ[54]. The ventral nerve cord (VNC) should be included since the circuits in the VNC are known to be crucial for behavior[55]. At least one optic lobe should be included to simplify analysis of visual inputs to the central brain. A whole brain connectome is preferable to the hemibrain, since then almost all cell types would have at least two examples, left and right, which would lend increased confidence to our reconstructions. It would also provide complete reconstruction to the many neurons that span the brain and are incomplete in the hemibrain. These three goals are combined in a project that is currently underway, reconstructing an entire male central nervous system (CNS) including the VNC and optic lobes.

We continue to improve sample preparation, imaging, and reconstruction to both to decrease the efforts expended on reconstruction and to allow reconstruction of more specimens. This includes multi-beam imaging, etching methods[56] that can handle larger areas, and yet better reconstruction techniques.

## 7 Author Contributions

ZL developed sample preparation and fixed and stained the sample; CSX, KJH, HFH developed the imaging hardware and imaged the sample; LS, ETT, DK, KAK, DA and SS aligned, flattened and integrated the slabs into a coherent volume; WK wrote and managed the versioned data system; SB, TZ, PH, LU, TD, DRS, DJO, NN, SP, JC, LS, ET, PS, TK wrote proofreading and analysis software; MJ, JMS, PHL, VJain developed and applied segmentation approaches; TB, LL, JMS developed analysis, visualization and pipeline software; MJ, LL, VJain developed and applied tissue classification approaches; GH developed and applied methods to identify synapses; KS, ST, JG, MI, TW, FL, KI, RP, JAH defined brain regions and cell types; JB and HO did the EM-optical mapping; CXA, DAB, SB, JAB, BSC NC, MC, MD, OD, BE, KF, SF, NF, AF, GPH, EMJ, SK, NAK, JK, SAL, AL, CM, EAM, SM, CM, MN, OO, NO, Co, NP, CP, TP, EEP, EMP, NR, CR, MKR, JTR, SMR, MS, AKS, ALS, AS CS, KS, NLS, MAS, AS, JS, ST, IT, DT, ET, TT, JJW AND TY performed proofreading; EN and CJK provided proofreading analytics; VJay, IAM, ST, KS, PR, RP, FL, GMR did biological interpretation and analysis; LS, SP did connectivity analysis and wrote the paper; JAH, FL, PR, RP managed proofreading; RG, GMR, SP managed the overall effort.

## Acknowledgments

We thank our colleagues at Janelia and the broader connectomics field for many helpful discussions and suggestions during the course of this work. We thank David Peale, Patrick Lee, and the Janelia Experimental Technology group for supporting the modifications of the FIB-SEM systems. Goran Ceric and other members of the Scientific Computing Systems and Scientific Computing Software Teams at Janelia provided critical support throughout this work. The Janelia Facilities group was essential in proving a stable environment for image collection. We thank Julia Buhmann and Jan Funke for help in implementing the synapse prediction algorithm described in [28]. Many colleagues at Janelia as well as Marta Costa, Greg Jefferis and others in Cambridge tested Neuprint performance prior to release. Major financial support for this work was provided by the Howard Hughes Medical Institute and Google Research. Feng Li, Philipp Schlegel and approximately 10 percent of the proofreading team were supported by a Wellcome Trust Collaborative Award (203261/Z/16/Z).

